# P2Y2 receptors are essential for hepatic clearance of uropathogenic *E. coli* in a murine sepsis model

**DOI:** 10.1101/2025.06.10.658777

**Authors:** Mette G. Christensen, Nanna Johnsen, Laura V. Sparsø, Thomas Corydon, Helle A. Praetorius

## Abstract

Urosepsis is a life-threatening condition most frequently caused by *E. coli* expressing important virulence factors, including α-haemolysin (HlyA). The pore-forming exotoxin HlyA releases ATP upon its insertion into cellular membranes, and the majority of the biological effects of HlyA are mediated through ATP-dependent P2-receptor activation, including the HlyA-mediated thrombocyte activation. We have recently shown that uropathogenic *E. coli* (UPEC) bind to thrombocytes immediately after entering the blood, and the following hepatic clearance leads to early thrombocytopenia during bacteraemia. Here, we demonstrate that P2Y_2_-deficient mice had markedly shorter survival (LD_50_ of 185 minutes) compared to wildtype (340 minutes), a response paralleled in mice infused with the P2Y_2_ receptor antagonist AR-C118925XX. The P2Y_2_^-/-^ mice exhibited a blunted sepsis-induced thrombocytopenia compared to wildtype and sepsis-induced reduction in mature neutrophils in the blood. Strikingly, the P2Y_2_-deficient mice had inadequate hepatic clearance of UPEC, resulting in the accumulation of bacteria in the lungs, while thrombocytes were mainly sequestered in the kidneys. Hence, it is likely that the insufficient hepatic elimination of UPEC is responsible for the reduced survival in the P2Y_2_^-/-^ mice. Taken together, we show that the lack of functional P2Y_2_ receptors is essential for fast and proper hepatic clearance of UPEC and the survival time during urosepsis. Moreover, the data support the notion that an early reduction in circulating thrombocytes is important for a relevant host response to acute bacteraemia.

## Introduction

Sepsis is a life-threatening condition defined by a dysregulated host response to infection. The urinary tract serves as the primary infection site in approximately 14% of all cases of sepsis [1], and patients with urological diseases and chronic kidney disease are more prone to contract urinary tract infections [2]. Generally, urinary tract infections (UTI) cover a spectrum from uncomplicated cystitis to more severe conditions like pyelonephritis and urosepsis, the latter exhibiting a 30-day mortality rate of up to 14% [3], and it is estimated that approximately 30% of all patients with pyelonephritis have bacteraemia [4]. The most frequent pathogen causing UTI as well as urosepsis is the Gram-negative bacterium *Escherichia coli* (*E. coli*) [5]. One of the central prognostic markers for an adverse outcome in sepsis is thrombocytopenia [6, 7], which has mainly been ascribed to the hypercoagulation state associated with sepsis. However, the immune modulatory effect of thrombocytes is receiving increasing attention, emphasising a role in the early response to infective agents in the blood (for review, see [8, 9]). We have recently shown that introducing uropathogenic *E. coli* (UPEC) into the blood is associated with an acute, marked reduction in circulating thrombocytes [10]. Interestingly, the UPEC immediately binds the thrombocytes for instant hepatic clearance [10], which most likely also involves mature neutrophils from the circulation. Remarkably, the liver Kupffer cells are circumvented in this process, whereas the sinus endothelial cells are shown to, at least transiently, bind complexes and UPEC in the clearance process [10]. This mechanism resembles what has been previously described for extravasations of metastatic melanoma (B16) cells, where binding of thrombocytes to the metastatic cancer cells facilitates neutrophil-dependent extravasation of the cancer cells [11].

Interestingly, this process requires endothelial expression of ATP-activated purinergic P2Y_2_ receptors [11]. The *E. coli* subtypes that cause pyelonephritis and urosepsis express various virulence factors [12, 13], including α-haemolysin (HlyA) [14]. In addition to being a virulence factor in ascending urinary tract infections (UTIs), HlyA increases mortality and is crucial for developing septic symptoms in mice with bacteraemia induced by *E. coli*, including the reduction in circulating thrombocytes [15]. HlyA is a pore-forming toxin that inserts itself receptor-independently into biological membranes, allowing ATP to be released directly through the pore into the extracellular environment [16]. The released ATP mediates the majority of HlyA’s biological effect via subsequent P2 receptor activation [17–19].

Based on this, we hypothesise that UPEC is cleared from the circulation through a mechanism similar to the one that drives the extravasation seen for cancer cells and, hence, that the P2Y_2_ receptor might be required for the hepatic clearance of UPEC. Thus, we here test the involvement of the P2Y_2_ receptor in UPEC clearance in a mouse model of urosepsis in the hope of advancing our understanding of urosepsis pathogenesis.

## Methods

### Animals

For the experiments, we used male, wildtype (WT) Balb/cJRj mice, either purchased from Janvier Labs (Saint-Berthevin, France) or bred at the Department of Biomedicine, Aarhus University. P2Y_2_-deficient (P2Y_2_^-/-^) mice on a Balb/cJRj background were also bred at the department after over 20 times backcross in Balb/cJRj, with new Balb/cJRj reintroduced regularly. All animals had free access to standard rodent chow and water, and the protocol has been approved by The Animal Experiment Inspectorate – Denmark (2020-15-0201-00422).

### Bacteria

The uropathogenic *Escherichia coli* (UPEC) strain ARD-6 (O6:K13:H1, Statens Serum Institut, Copenhagen, Denmark) was used as previously described [20–22] or transfected with a GFP-encoding plasmid with ampicillin restriction (ARD-6/EGFP-pBAD)[23]. Stocks of UPEC (15% glycerol Lysogeny Broth (LB)-media) were kept at - 80°C. New colonies were prepared with 1 µl of UPEC stock streaked out on an LB-agar plate (100 μg ml^-1^ ampicillin for ARD-6/EGFP-pBAD) for incubation overnight (37°C) prior to storage at 4°C for up to one month. For overnight cultures, a single colony was transferred to 4 ml LB-medium (containing 100 μg ml^-1^ ampicillin for ARD-6/EGFP-pBAD) and incubated aerobic at 37°C, 250 rpm. The overnight cultures were centrifuged (10 minutes, 1300g) and washed (5 minutes, 1300g), resuspended in sterile saline and counted by flow cytometry (BD Accuri 6C+, BD biosciences, New Jersey, USA). ARD-6/EGFP-pBAD were prior to washing centrifuged (10 minutes, 1300 g) and incubated for 3 hours in LB-medium containing 100 μg ml^-1^ ampicillin/0.2% w/v L-arabinose.

### Induction of sepsis in mice

Mice (8-10 weeks, 24.6 ± 0.1 g) were anaesthetised with a subcutaneous injection of ketamine (100 mg kg^-1^) and xylazine (10 mg kg^-1^) and placed on a heating plate (38°C). For the P2Y_2_ antagonist experiments, mice received a bolus injection of AR-C118925XX (0.37 μg g^-1^) or vehicle (DMSO, 10.6 μl ml^-1^) before continuous infusion with AR-C118925XX (2.03 μg hour^-1^) or vehicle throughout the experiment. The mice were infused (66 μl hour^-1^) via one of the tail veins (27G needle, Masterflex multi-syringe pump, Cole-Parmer, IL, USA). UPEC (330 10^6^ or 41 10^6^) were injected into the tail vein in pure, sterile saline (WT and P2Y_2_^-/-^) or for P2Y_2_ receptor antagonist with (AR-C118925XX) or vehicle added to the solution. For survival data, the mice were observed for up to six hours, and the time of events was noted, whereas, for the other experiments, the mice were terminated at various time points after the collection of blood from the inferior vena cava with a citrate-containing syringe.

### Quantification of circulating bacterial load

Whole blood (10 μl) from mice exposed to UPEC or vehicle was isolated for blood culture. The sample was diluted 1:500 with sterile saline (0.9%), and 50 μl from this preparation was streaked on an LB agar plate and incubated overnight at 37°C for determination of the number of colony-forming units (CFU). For flow cytometric UPEC count, fixed blood samples were diluted 1:5.2 and incubated for 30 minutes with 3.83 µg ml^-1^ anti-CD42d-APC (eBioscience™, Invitrogen) in the dark with a following dilution of 1:76 in PBS. A compensation matrix was generated from single staining and median fluorescence values. Thrombocytes (APC-positive), UPEC (GFP-positive), and their aggregates were quantified via flow cytometry (BD Accuri 6C, BD Biosciences). See [10] for gating strategies.

### Detection of leukocytes

The number of thrombocytes, neutrophils, monocytes and platelet-leukocyte aggregates was determined in whole blood by flow cytometry (BD Accuri C6, BD Biosciences). To detect leukocytes and platelets, whole blood diluted (1:13) in Ca^2+^-free buffered salt solution with apyrase (0.03 U ml^-1^) was incubated for 30 minutes in the dark either with the neutrophil marker anti-Ly6G-FITC (23.3 mg ml^-1^, eBioscience^TM^, Invitrogen) and anti-CD42d-APC (4.65 mg ml^-1^, eBioscience^TM^, Invitrogen) for detection of thrombocytes, or with the combination of anti-Ly6C-FITC (4.57 mg ml^-1^, Abcam) and anti-CD11b-PE-Cy7 (3.65 mg ml^-1^, eBioscience^TM^, Invitrogen) as monocyte marker with the thrombocyte marker anti-CD42d-APC (4.65 mg ml^-1^, eBioscience^TM^, Invitrogen). Before analysis, the samples were diluted 1:76 with fixative (0.02% formaldehyde in PBS). The preparation was analysed by the BD Accuri C6 analysis software (BD Biosciences, NJ, USA); monocyte-platelet aggregations were determined as Ly6C^+^, CD11b^+^, CD42d^+^ events and neutrophil-platelet aggregations as Ly6G^+^, CD42d^+^ events (see suppl. fig. S1 and S2). Neutrophils were gated for maturity based on the maturity anti-Ly6G-FITC signal (see [10]).

### Measurement of intravascular haemolysis

Blood samples were centrifuged for 10 minutes at 1162*g,* and plasma was diluted 1:8 with saline before measurement of the absorbance at 410 nm on a spectrophotometer (Ultraspec III, LKB, Biochom). The remaining plasma was stored at -80°C for later analysis of cytokines, thrombin-antithrombin (TAT)-complexes and platelet factor 4 (PF-4).

### Measurement of IL-6, KC, TNF-α and IL-1β

Levels of IL-6, keratinocyte chemoattractant (KC, murine equivalent to human IL-8), TNF-α and IL-1β were measured using a cytometric bead array flex set (BD Bioscience, NJ, USA) according to the manufacturer’s instructions and analysed by flow cytometry (BD Accuri C6, BD Biosciences). Cytokines were measured in plasma stored at -80°C for up to 3 months before analysis.

### Markers for coagulation and platelet activation

Thrombin-antithrombin (TAT) complexes were determined in plasma samples using a TAT-complex mouse ELISA kit (Abcam, Cambridge, UK) according to the manufacturer’s instructions. PF-4 were determined in plasma using a mouse Platelet Factor 4 CXCL4 ELISA kit (Sigma-Aldrich, St. Louis, MO, USA) according to the manufacturer’s protocol. Plasma samples were stored for up to 3 months at -80°C before TAT complex and PF-4 measurements.

### Statistics

Statistical analyses were performed using GraphPad Prism v. 9.0. Survival and haemoglobinuria were analysed by Kaplan-Meier plots and the Log-rank (Mantel-Cox) test. The remaining data were tested for normal distribution using the Kolmogorov-Smirnov test. Normally distributed data were tested by two-way Anova a Šidák multiple comparisons, and data which were not normally distributed were analysed with the Kruskal-Wallis test followed by Dunn’s multiple comparisons test to compare either between genotypes, treatment or sepsis/non-sepsis. All data are presented as mean ± standard error of the mean (SEM). Data were considered statistically significantly different when the p-value was less than 0.05.

## Results

### Lack of functional P2Y_2_ receptor reduces survival of UPEC-induced sepsis in mice

We have previously demonstrated that UPEC binds to thrombocytes immediately after being introduced into the blood and that bacterial complexes are rapidly cleared by the liver [10]. The P2Y_2_ receptor is essential for thrombocyte-dependent extravasation of cancer cells during metastasis formation [11], and we speculated that the elimination of UPEC might occur through a similar mechanism. Therefore, we investigated if P2Y_2_ receptors are involved in hepatic UPEC clearance. We used an established model for urosepsis in mice, where the bacterial load was titrated to result in an LD_50_ of around 4 hours. Figure 1A shows the survival of WT and P2Y_2_^-/-^ mice exposed to 3.3 10^8^ UPEC or saline as a bolus injection in a tail vein. Mice exposed to saline survived for the entire observation period except for one P2Y_2_^-/-^ the mouse that died immediately before the termination of the experiment. In exposure to UPEC, the P2Y_2_^-/-^ dies statistically significantly earlier (median survival of 185 minutes) than the WT (median survival of 340 minutes). This finding was confirmed in WT mice continuously infused with the P2Y_2_ receptor antagonist AR-C118925XX (2.03 μg h^-1^, Fig. 1B) compared to vehicle-infused mice. Mice infused with AR-C118925XX died statistically significantly earlier than the vehicle controls when exposed to 3.3 10^8^ UPEC, whereas the saline-infused mice survived for the entire observation period.

**Figure 1.**
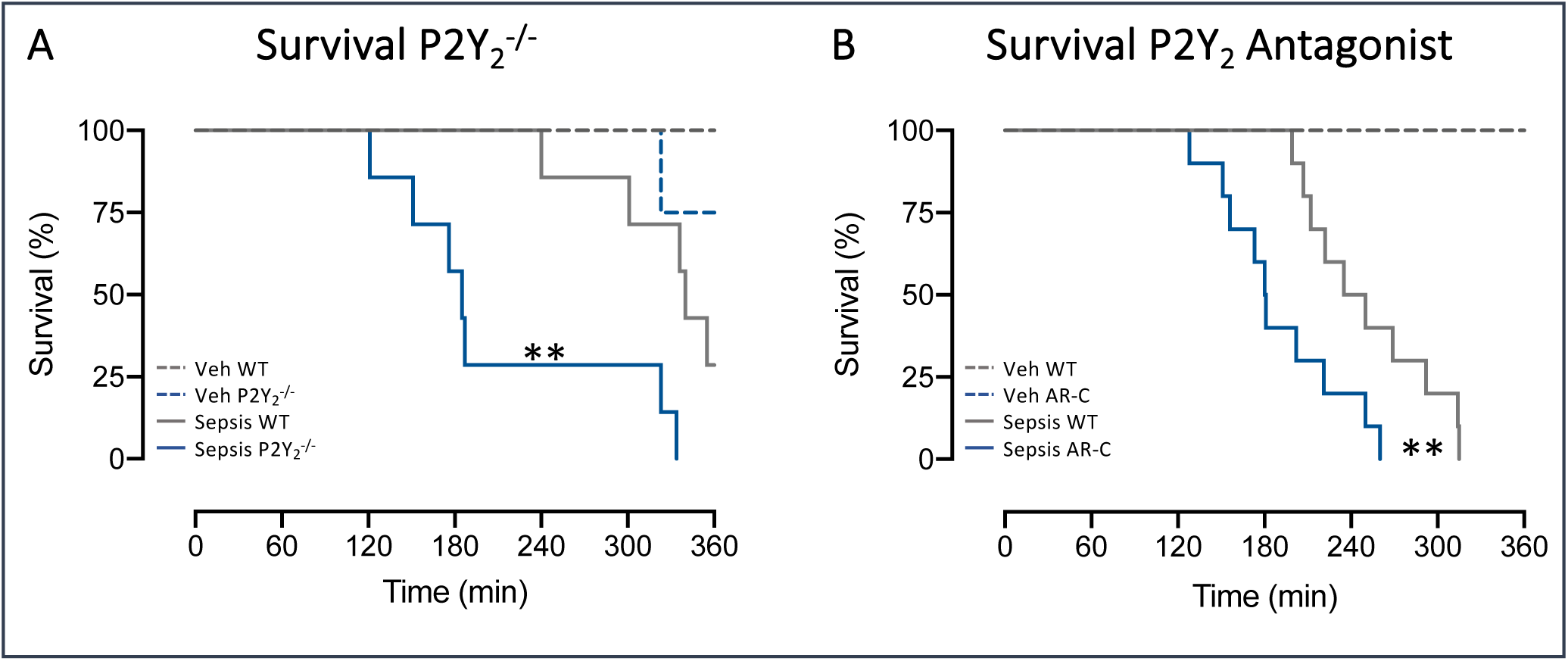
Lack of inhibition of the P2Y_2_ receptors increases mortality in mice with UPEC-induced sepsis. (**A**) Shows the survival in WT (*n*=7) and P2Y ^-/-^ (*n*=7) mice injected with 3.3 10^8^ UPEC. The dotted lines represent control injection with saline in WT (*n*=4) and P2Y ^-/-^ (*n*=4) mice. (**B**) illustrating the survival of WT mice infused with the P2Y receptor antagonist AR-C118925XX (AR-C, 2.03 μg hour^-1^, *n*=10) or vehicle (*n*=10) after injection with 3.3 10^8^ UPEC. The dotted lines represent mice injected with saline (n=3 for both vehicle and AR-C treated animals). Data are shown as Kaplan-Meier plots, and ** indicates *p*<0.01

The discrepancy in survival between WT mice and P2Y_2_^-/-^ mice cannot be immediately explained by a difference in the number of circulating UPEC (Fig. 2A). The figure shows that UPEC is cleared from the circulation within the first hour of the infection both in the WT and the KO mice, although the scatter in the data is more extensive in the KO mice. The systemic response is seemingly developed in parallel in the two mouse types with similar onset of haemoglobinuria and an increase in proinflammatory cytokines, although only the P2Y_2_^-/-^ mice showed an increase in IL-1β in the observation period (Fig. 2B-F). A somewhat similar pattern was observed with an infusion of AR-C118925XX compared to a vehicle infusion. However, Figure 3A reveals a higher level of circulating UPEC in mice exposed to AR-C118925XX infusion compared to vehicle-infused controls immediately after injection of UPEC. Please note that time zero represents blood sampling after systemic distribution of UPEC (ca. 10, ∼75 heartbeats). The mice exposed to AR-C118925XX get haemoglobinuria earlier than the vehicle controls (Fig. 3B), but the profile with regard to proinflammatory markers is reasonably similar in the two groups; only keratinocyte chemoattractant, binding to murine IL-8 receptors selectively, increases in the vehicle-infused mice, without becoming statistically significantly higher in the AR-C118925XX-infused (Fig. 3C-F).

**Figure 2.**
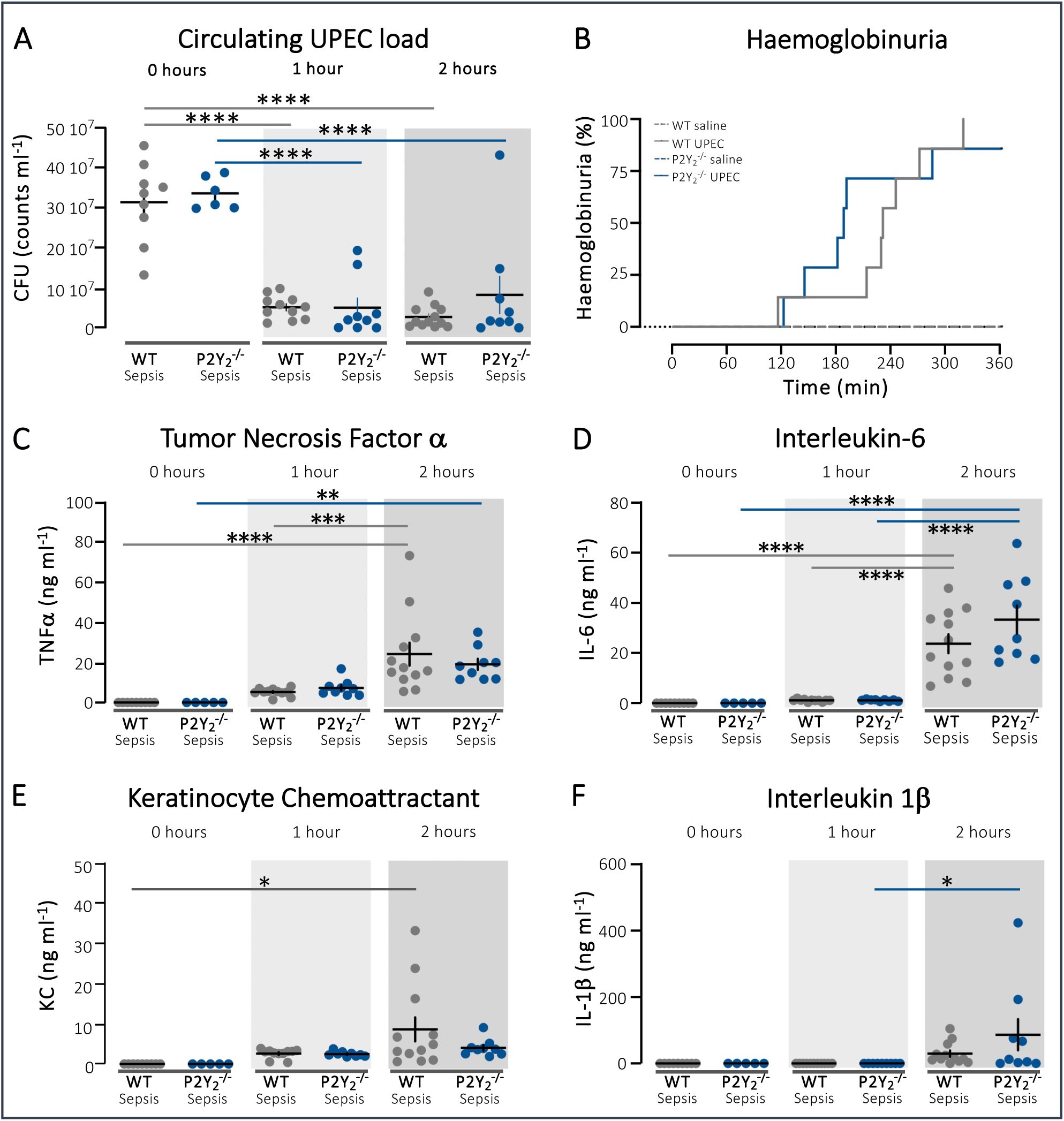
Bacterial load and systemic response to sepsis induced with UPEC in WT and P2Y ^-/-^. (**A**) shows the bacterial load as colony forming units (CFU) in blood from mice injected with 3.3 10^8^ at time 0 (*n*=9 for WT and *n*=5 for P2Y ^-/-^) and 2 hours (*n*=12 for P2Y ^+/+^ and *n*=9 for P2Y ^-/-^). (**B**) depicts the development of haemoglobinuria in the mice as a measure of the systemic response to circulating UPEC. (C-F) show the corresponding change in pro-inflammatory cytokines (TNF-α,IL-6, KC and IL-1β) in WT and P2Y ^-/-^ mice with sepsis with 3.3 10^8^ UPEC (*n*=9-12 for WT and *n*=5-9 for P2Y ^-/-^ sepsis). Data are shown as single observation and mean±SEM, * indicates *p*<0.05, \*\**p*<0.01, \*\*\**p*<0.001 and \*\*\*\**p*<0.0001.

**Figure 3.**
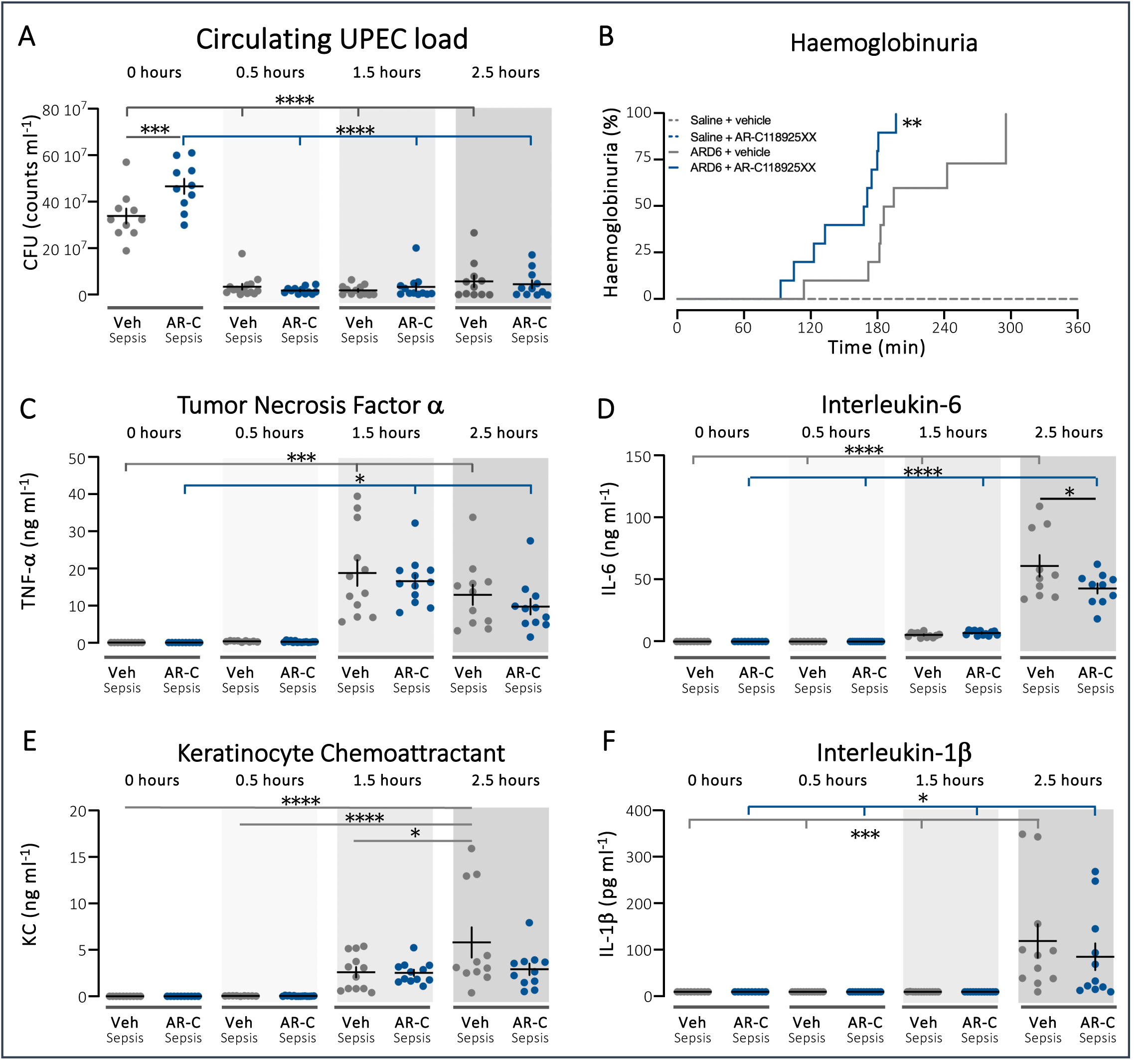
Bacterial load and systemic response to sepsis induced with UPEC in WT infused with the P2Y_2_-antagonist AR-C118925XX (AC-R, 2.03 μg h^-1^) or vehicle infusion. (**A**) shows the bacterial load as CFU in blood from WT mice with sepsis at time 0-2.5 (*n*=10-14) for both groups after induction of sepsis with 3.3 10^8^ UPEC. (**B**) illustrates the development of haemoglobinuria in the mice as a measure of the systemic response to circulating UPEC. (**C-F**) show the corresponding change in pro-inflammatory cytokines (TNF-α,IL-6, KC and IL-1β) in WT mice exposed to AR-C118925XX or vehicle after induction of sepsis with 3.3 10^8^ UPEC. Data are presented as single observations with mean±SEM, * indicates *p*<0.05, *** *p*<0.001 and \*\*\*\**p*<0.0001.

### The sepsis-induced reduction in circulating thrombocytes is delayed in P2Y_2_^-/-^ mice

We have previously shown that the introduction of UPEC in circulation markedly reduces the number of thrombocytes in the blood as a result of the hepatic clearance of UPEC/thrombocyte complexes [10]. Figure 4A shows that after injection of 3.3 10^8^ UPEC, we observe a reduction in circulating thrombocytes in WT and P2Y_2_^-/-^ mice over time. However, the reduction in thrombocyte count becomes significantly different in the P2Y_2_^-/-^ after 2 hours, whereas the thrombocyte number is already significantly reduced after one hour in the WT mice. This occurs despite platelet factor 4 being significantly elevated in the P2Y_2_^-/-^ mice, whereas this is only a trend in the WT mice (Fig. 4B). Notably, we do not observe increased coagulation in either of the mice types, although there is a tendency for an increase in thrombin-antithrombin complexes 2 hours after UPEC injection. To further substantiate whether there is a reduced clearance/tissue adherence of thrombocytes in the P2Y_2_^-/-^ mice, we examined the reduction in circulating thrombocytes with a lower bacterial load. Figure 4D shows a 31% reduction in circulating thrombocytes 2.5 hours after injection of 4.1 10^7^ UPEC compared to saline controls (*p*=0.0002). Interestingly, the sepsis-induced reduction of thrombocytes was not detectable in P2Y_2_^-/-^ mice (p=0.481), and thrombocyte count during sepsis was statistically significantly higher in the P2Y_2_^-/-^ mice compared to WT mice with sepsis (Fig. 4D). In parallel, we detected an increase in TAT complexes in the WT mice 2.5 hours after exposure to 4.1 10^7^ (p=0.027), whereas this was not the case in the P2Y_2_^-/-^ mice (Suppl. Fig. 1). This relatively subtle phenotype was not echoed in the experiments with the P2Y_2_ agonist AR-C118925XX (Suppl. Fig. 2). To evaluate the effect of the P2Y_2_ receptor in the clearance of UPEC-thrombocytes complexes in the blood, we monitored the removal of bacteria in a period up to 10 minutes after injection in mice. Figure 5A shows that both WT and P2Y_2_^-/-^ mice could clear the bacteria fast, whereas the fall in total circulating thrombocytes is not statistically significant yet in any of the mouse strains (Fig. 5B). The UPEC bacteria complexes are cleared in both phenotypes, with a larger deviation in the P2Y_2_^-/-^ at time 5 minutes compared to the WT mice. This picture is paralleled when infusing AR-C118925XX compared to vehicle infusion (Fig. 5D-F). Total UPEC and UPEC-complexes are fast cleared irrespective of infusion type. However, the UPEC-induced thrombocyte reduction is slightly delayed in AR-C118925XX infused mice compared to vehicle.

**Figure 4.**
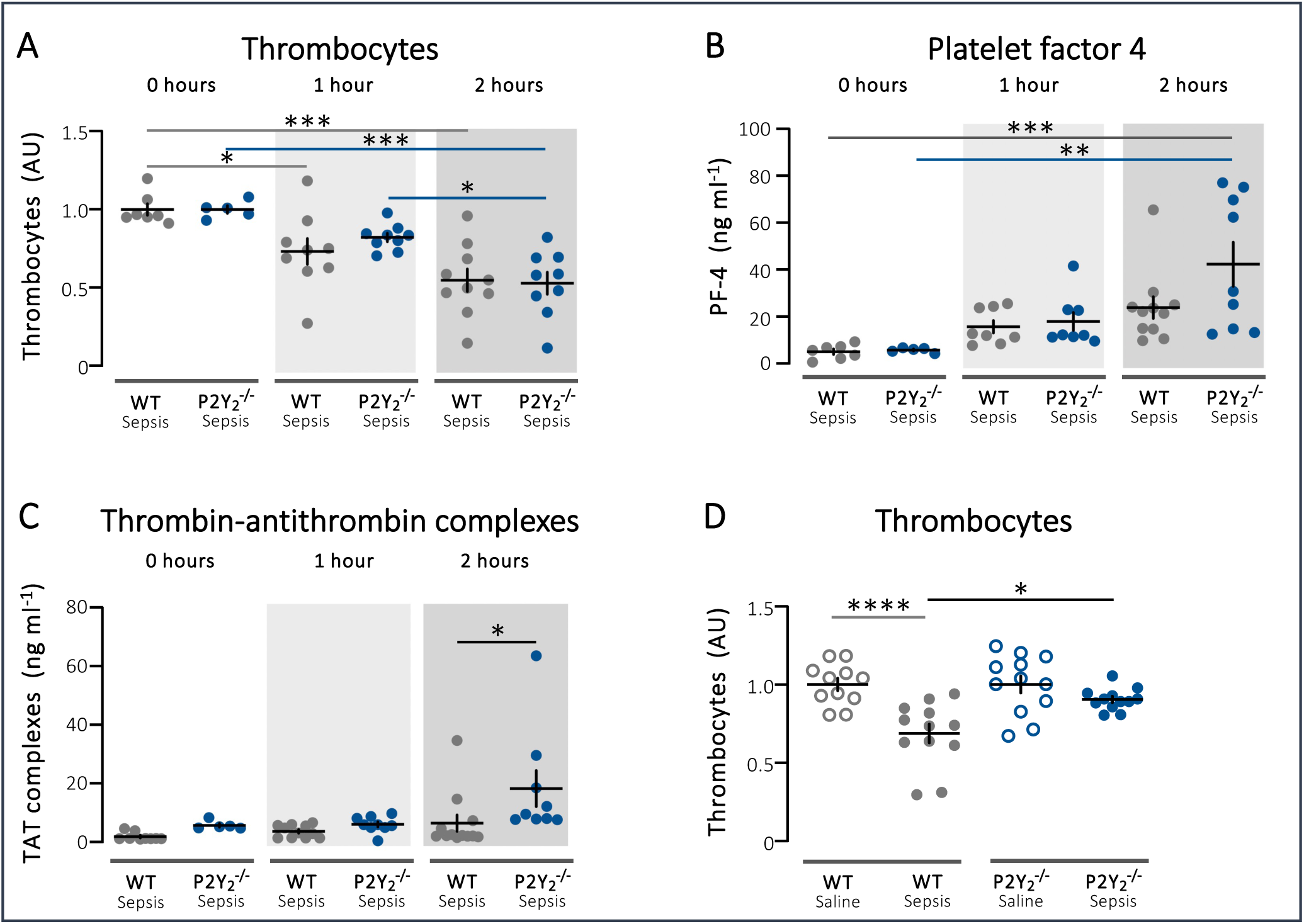
The sepsis-induced fall in circulating thrombocytes is delayed in P2Y ^-/-^ mice with UPEC-induced sepsis. (**A**) shows the relative number of thrombocytes in WT and P2Y ^-/-^ mice from 0-2 h after injection of 3.3 10^8^ UPEC (*n*=5-10 in the groups), (**B**) illustrates the corresponding change in thrombocyte activation marker platelet factor-4 (PF-4) in WT and P2Y ^-/-^ in plasma of the same animals. (**C**) Shows the change in intravascular formation of thrombin-antithrombin (TAT) complexes in plasma in the same animals as in A. (**D**) depicts the relative change in number of thrombocytes in WT and P2Y ^-/-^ mice 2.5 hours after injection of either saline or 4.1 10^7^ UPEC. (*n*=11-12 in each group). Data are given as mean±SEM, * indicates *p*<0.05, \*\**p*<0.01 and \*\*\*\**p*<0.0001.

**Figure 5.**
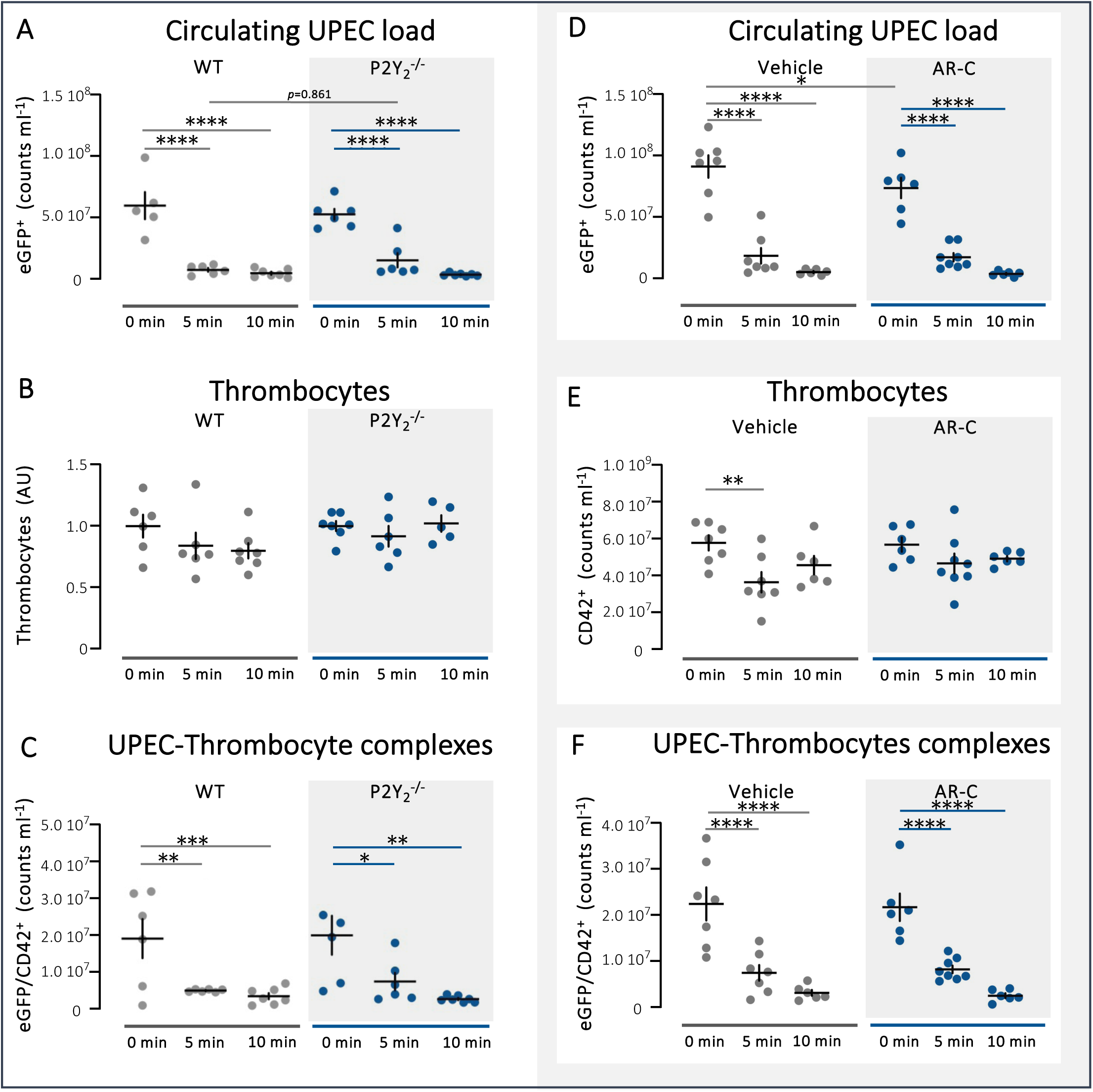
The effect of the P2Y_2_ receptor on the clearance of UPEC-thrombocyte complexes. (**A**) shows UPEC count in the blood of WT mice at time zero, 5 min and 10 min after injection of 3.3 10^8^ UPEC (*n*=6-7) (**B**) shows the corresponding thrombocyte count and (**C**) the UPEC-thrombocyte complexes. (**D**) shows UPEC count in the blood of P2Y ^-/-^ mice at time zero, 5 min and 10 min after injection of 3.3 10^8^ UPEC (*n*=6-7) (**E**) shows the corresponding thrombocyte count and (**F**) the UPEC-thrombocyte complexes. All data are given as mean±SEM, * indicates *p*<0.05, ** *p*<0.01 and \*\*\*\**p*<0.0001.

### Recruitment of immune cells does not require functional P2Y_2_ receptors

P2Y_2_ receptors are also expressed on a number of immune cells, and thus, one could speculate that the increased mortality, seen in both P2Y_2_^-/-^ mice and mice treated with a P2Y_2_ receptor antagonist, might result from impaired recruitment of immune cells or immune dysfunction. As shown in Figure 5A, the number of circulating neutrophils does not result in a statistically significant change during the first two hours of infection, either in the WT or P2Y_2_^-/-^ mice (Fig. 6A). Similarly, the number of neutrophil/thrombocyte complexes does not change and are similar in WT or P2Y_2_^-/-^ mice over time (Fig. 6B). The stable number of neutrophils hides an early fall in mature neutrophils within the first hour of infection, with a tendency towards an increase in immature neutrophils in WT animals (Fig. 6C). Interestingly, in the P2Y_2_^-/-^ mice, the tendency for the mature neutrophils to fall in the early phases of sepsis did not become statistically significant. However, we could clearly see the recruitment of immature neutrophils (Fig. 6D, *p*=0.0181 for comparison between 0 and 2 hours). We did not observe any significant changes in the number of circulating monocytes (Fig. 6E) or monocyte-thrombocyte complex formation (Fig. 6F).

**Figure 6.**
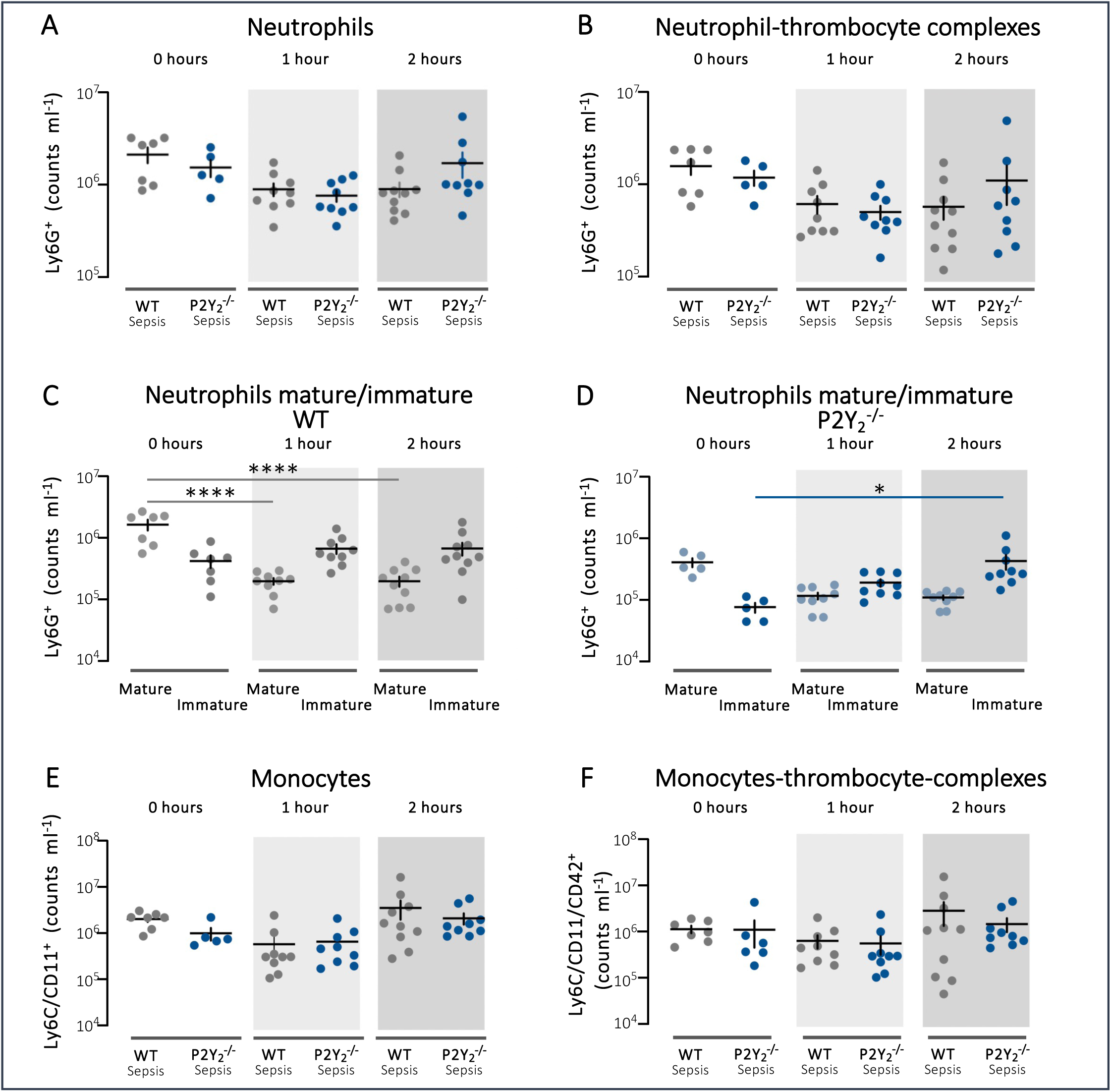
The sepsis-induced changes in circulating neutrophils and neutrophil-thrombocyte complexes. (**A**) shows the neutrophil count in blood from WT and P2Y ^-/-^ mice after injection of 3.3 10^8^ UPEC (*n*=5-10) (**B**) shows the corresponding formation of neutrophil-thrombocyte complexes. (**C**) illustrates the change in mature and immature neutrophils over time in WT mice exposed to 3.3 10^8^ UPEC or saline, whereas (**D**) shows the corresponding data in the P2Y ^-/-^ mice. (**E**) depicts the monocyte count in the blood in the same animals and (F) monocyte-thrombocyte complexes. All data are given as mean±SEM, * indicates *p*<0.05, \*\*\*\**p*<0.0001.

### P2Y_2_ receptors are essential for specific hepatic clearance of UPEC-thrombocyte complexes

We have previously shown that UPEC, alone or in complex with thrombocytes, is effectively and rapidly cleared in the liver [10]. Figure 7A confirms this and shows a marked accumulation of UPEC in the liver 30 minutes after injection of 3.3 10^8^ UPEC into the blood in WT mice, with very few escaping for tissue accumulation elsewhere. Similarly, we replicated the substantial accumulation of thrombocytes and UPEC-thrombocyte complexes in the liver within the first 30 minutes of exposure to UPEC (Fig. 7B-C). It must be noted that thrombocytes are also found in substantial amounts in the spleen, and our previous time-lapse study showed that the fraction of thrombocytes in the spleen is constant over time; there is not an increase during sepsis [10].

**Figure 7.**
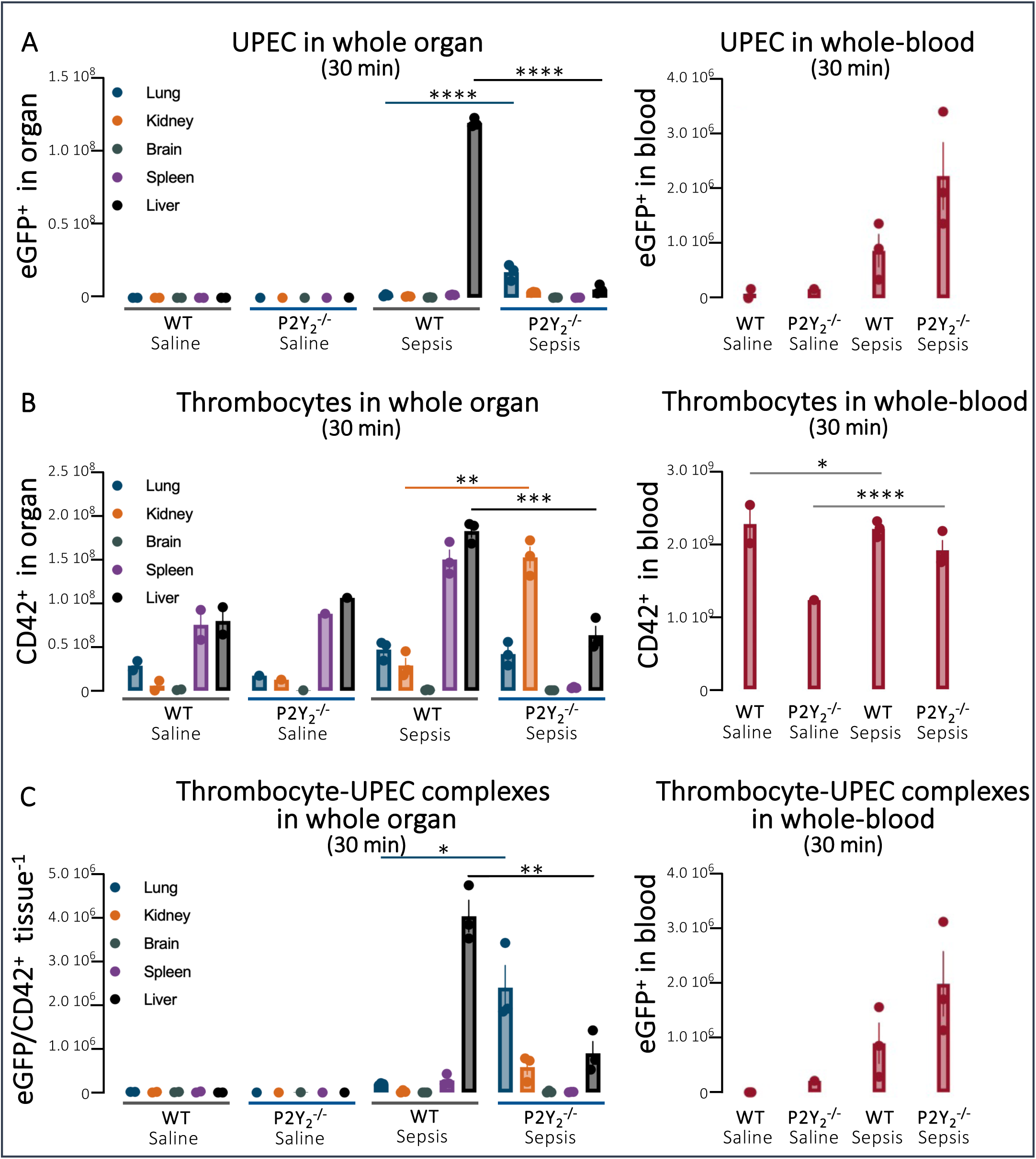
Tissue-specific accumulation of UPEC and thrombocytes. (**A**) shows the tissue-dependent clearance of UPEC in WT and P2Y ^-/-^ mice 30 minutes after injection of 3.3 10^8^ ARD-6/EGFP-pBAD, and (**B**) the corresponding tissue accumulation of thrombocytes and (**C**) UPEC-thrombocyte complexes. Data are given as mean±SEM, and * indicates *p*<0.05, \*\**p*< 0.01, \*\*\**p*<0.001 and \*\*\*\**p*<0.0001.

In contrast, P2Y_2_^-/-^ mice show quite a different pattern, with only a minor fraction of the UPEC being captured in the liver (Fig. 7A), but are now passing on to the lungs and, to a lesser extent, the kidneys. Similarly, the thrombocytes are also less likely to accumulate in the liver of the P2Y_2_^-/-^ mice and are in this phenotype accumulating in the kidneys (Fig. 7B). Likewise, the UPEC-thrombocyte complexes pass the liver and accumulate in the lungs (Fig. 7C). These data show that the P2Y_2_^-/-^ mice lack the effective hepatic clearance system for UPEC-thrombocyte complexes. Interestingly, thrombocytes preferentially accumulate in the kidney, and the UPEC are primarily trapped in the lung in this transgenic mouse, illustrating that the accumulation no longer depends on complex formation between the two. Moreover, we observe that the splenic thrombocyte reservoir of thrombocytes is drained in the P2Y_2_^-/-^ mice during the first 30 minutes of infection, likely contributing to the protracted development of thrombocytopenia in these mice. A similar pattern of clearance is seen when the data are given, not as whole-organ but per gram tissue (Suppl. Fig. 5). We speculate that this altered pattern in UPEC and thrombocyte clearance explains both the delayed reduction in thrombocytes, the tendency towards dampened clearance of UPEC and mature neutrophils, and the reduced survival in P2Y_2_^-/-^ mice.

## DISCUSSION

Urosepsis is a life-threatening systemic reaction to bacteraemia, with uropathogenic bacteria entering the blood from the urinary tract. Admissions under the diagnosis of sepsis are rising [24], frequently with the urinary tract as the primary infection site [25, 26]. In the urinary tract, the bacteria are positively selected for virulence, allowing them to survive, colonise, and, via the well-vascularised kidney, disseminate into the blood. We have previously shown that UPEC instantaneously binds to thrombocytes upon entering the blood and that a following hepatic clearance may result in an early reduction in circulating thrombocytes, depending on the bacterial load [10]. α-Haemolysin (HlyA) is a key virulence factor in both the development of UTIs and even during the progression of urosepsis caused by HlyA-producing *E. coli* [27]. As an exotoxin, HlyA inflicts its actions on the host cells both through the direct effect of membrane pore-formation and through the release of ATP, which, as a paracrine factor, amplifies the host cell response [17, 21, 22] and markedly aggravates the reduction in circulating thrombocytes in response to *E. coli* in the circulation [27].

Here, we show that mice deficient in the ATP-sensitive P2Y_2_ receptor exhibit reduced survival time during sepsis induced by UPEC injection, a finding underscored by a substantial reduction of survival time after infusion of the P2Y_2_ antagonist AR-C118925XX in the same model. This distinct phenotype is not equally obvious when looking at the blood parameters, although there is a tendency towards a less efficient bacterial clearance and a higher level of IL-1β in the P2Y_2_^-/-^ mice and a lower initial clearance of bacteria after infusion of AR-C118925XX. There was no difference in the ability of the UPEC to bind to the thrombocytes between the two phenotypes, but interestingly, the P2Y_2_^-/-^ mice showed delayed clearance of thrombocytes in response to UPEC-induced bacteraemia. We have previously demonstrated that injection of UPEC inflicts a rapid and substantial reduction in the circulating thrombocytes, which depends on bacterial load [10]. Hence, based on the survival data alone, one would have expected the thrombocytes to be markedly lower in the P2Y_2_^-/-^ mice compared to WT mice. Although we did observe a reduction in circulating thrombocytes in response to UPEC in the P2Y_2_^-/-^ mice, the reduction occurred with a delay despite a tendency for a higher degree of thrombocyte activation measured as PF-4 release. By reducing the dose of UPEC eight times, this effect became even more apparent, and we could only observe the sepsis-induced reduction of circulating thrombocytes in the WT mice and not in the P2Y_2_^-/-^ mice. In this context, it must be noted that thrombocytes, although very responsive to ATP and its degradation products, do not express the P2Y_2_ receptor (for an overview, see [28]), and hence, the reduced clearance is secondary to reduced thrombocyte activation. This is further supported by the data on PF-4 release and thrombin-antithrombin complex formation in WT and P2Y_2_^-/-^ mice.

In our previous study of hepatic bacterial clearance of UPEC-thrombocyte complexes, we observed a corresponding reduction in total circulating neutrophils. In this study, this was not statistically significant either in WT or in P2Y_2_^-/-^ mice. However, we found a significant early reduction in mature neutrophils in WT mice. The P2Y_2_^-/-^ mice showed the same tendency, but this did not become statistically significant. These data support the earlier proposed notion that neutrophils may be required for the hepatic clearance of circulating UPEC. The data also confirm our previous finding that the acute reduction in thrombocytes in response to UPEC is not caused by increased neutrophil-thrombocyte or monocyte-thrombocyte complex formation, although this type of complex formation was previously shown to be elevated during sepsis [10, 29–31]. In addition, the data shows that the reduced median survival time in P2Y_2_^-/-^ is not explained by a decreased neutrophil response.

Strikingly, the organ distribution of UPEC, thrombocytes and their complexes showed a completely different pattern in the P2Y_2_^-/-^ mice compared with WT. In the WT, we confirm a marked accumulation of UPEC and thrombocytes, of which some are still in complexes, in the liver within the first 30 minutes after injection of UPEC. In the P2Y_2_^-/-^ mice, very few UPEC are accumulated in the liver. Instead, they build up in the lungs and, to a lower degree, the kidneys. In the P2Y_2_^-/-^ mice, the thrombocyte accumulation seemingly does not follow UPEC to the same degree as in the WT. In the P2Y_2_^-/-^ mice, the thrombocytes mainly accumulate in the kidneys and only a minor proportion in the lungs. This suggests that the thrombocytes and UPEC are only, to a minor degree, removed from the circulation in the complex. Moreover, the pool of thrombocytes in the spleen has been mobilised in the P2Y_2_^-/-^ mice, which may indicate an increased need for acute buffering of free UPEC in the P2Y_2_^-/-^ mice during sepsis.

Our previous study on UPEC clearance in the liver revealed very little interaction with the hepatic Kupffer cells. This was surprising as Kupffer cells are generally considered to be responsible for the clearance of larger infectious agents (<1 µm), where the smaller particles (<0.3µm) are cleared by the sinusoidal endothelial cells (for review, see[32]). However, using flow cytometry, we found substantially increased interaction with endothelial cells, potentially suggesting that UPEC, via complex formation with thrombocytes, might circumvent adhesion to Kupffer cells. The current study further supports this notion as the P2Y_2_ receptor has been demonstrated to have a functionally relevant expression in endothelial cells (for review, see [33]) that is completely absent in P2Y_2_^-/-^ C57BL/6 background [34, 35] and also in P2Y_2_^-/-^ Balb/cJ background [36]. Cell type-specific RNAseq and proteomics on hepatic cells fail to show expression of P2Y_2_ receptors in Kupffer cells in mice [37] or humans [38], which is known to have P2X_7_ and P2Y_6_ as their primary purinergic receptors [39]. It must be noted that the RNAseq did not find P2Y_2_ in the sinusoidal endothelial cells either; however, they have been detected functionally in these cells in other studies [40]. Moreover, our data suggests that UPEC could be cleared in the liver by a very similar system as reported for cancer cells via thrombocyte binding and neutrophil-depended transport across the endothelium in a P2Y_2_ receptor-dependent fashion [11], a mechanism that has recently been shown to be relevant for liver metastasis via gap-formation in the sinusoidal endothelial cells [41]. Although the evidence is circumstantial, it is most likely that a lack of-P2Y_2_ on the endothelium results in the P2Y_2_^-/-^ mice’ insufficient hepatic clearance of UPEC. Since the main difference lies in the tissue distribution and not in the overall recruitment of neutrophils and proinflammatory response, it is likely that the phenotype is mainly endothelial-dependent, although this would have to be verified experimentally.

Considering the literature, it is surprising that we find the P2Y_2_^-/-^ mice to be more susceptible to urosepsis. An earlier study from Inoue et al. shows a marginally longer survival of P2Y_2_^-/-^ mice compared to controls after caecal ligation and puncture (CLP) [42]. However, the CLP model tests both the dissemination of local mixed infection and the response to bacteria in the blood, which is not immediately comparable to our model. Moreover, the hepatic clearance system for uropathogenic *E. coli* is likely to be specific to capsule-carrying Gram-negative bacteria that are relatively protected against complement-induced hepatic clearance [43], a feature also seen for Kupffer cell detection of *Klebsiella pneumoniae* [44] and *Streptococcus pneumoniae* [45]. Since the agent that causes the sepsis after caecal ligation is unknown, it may potentially be handled very differently and hence not be removed by the P2Y_2_ receptor-dependent clearance system.

In conclusion, the P2Y_2_ receptor is exceedingly important for normal host response to invading bacteria, and the lack of a P2Y_2_ receptor prohibits the normal hepatic clearance of invading UPEC. Moreover, the study underscores the essential function of thrombocytes in the early response to invading bacteria.

## Acknowledgement

We would like to thank Helle Jakobsen for skilled technical support. This study was funded by the Novo Nordisk Foundation (NFF 0064953).

---

**Supplementary Figure 1.**
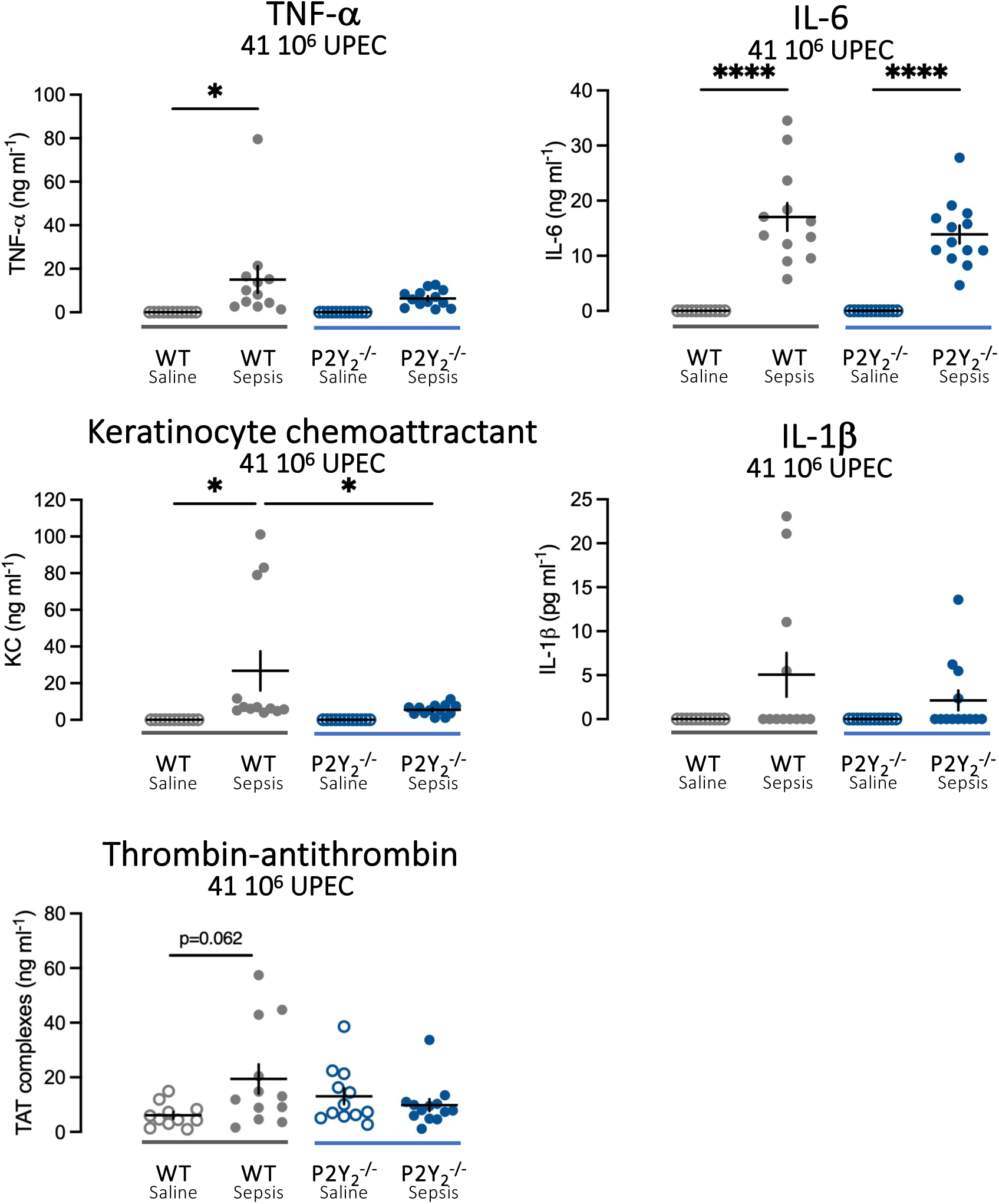
Change in proinflammatory cytokines and thrombin-antithrombin complexes in response to UPEC (41 106 ) after 2.5 hours in WT and P2Y2 -/-. (**A**) TNFa, (**B**) IL-6 (**C**) Kertinocyte Chemoattractant (KC) and (**D**) IL-1b. (**E**) shows the corresponding changes in the thrombinantithrombin complex. Data are shown as single observation and mean±SEM., * indicates p<0.05 and ****p<0.0001.

**Supplementary Figure 2.**
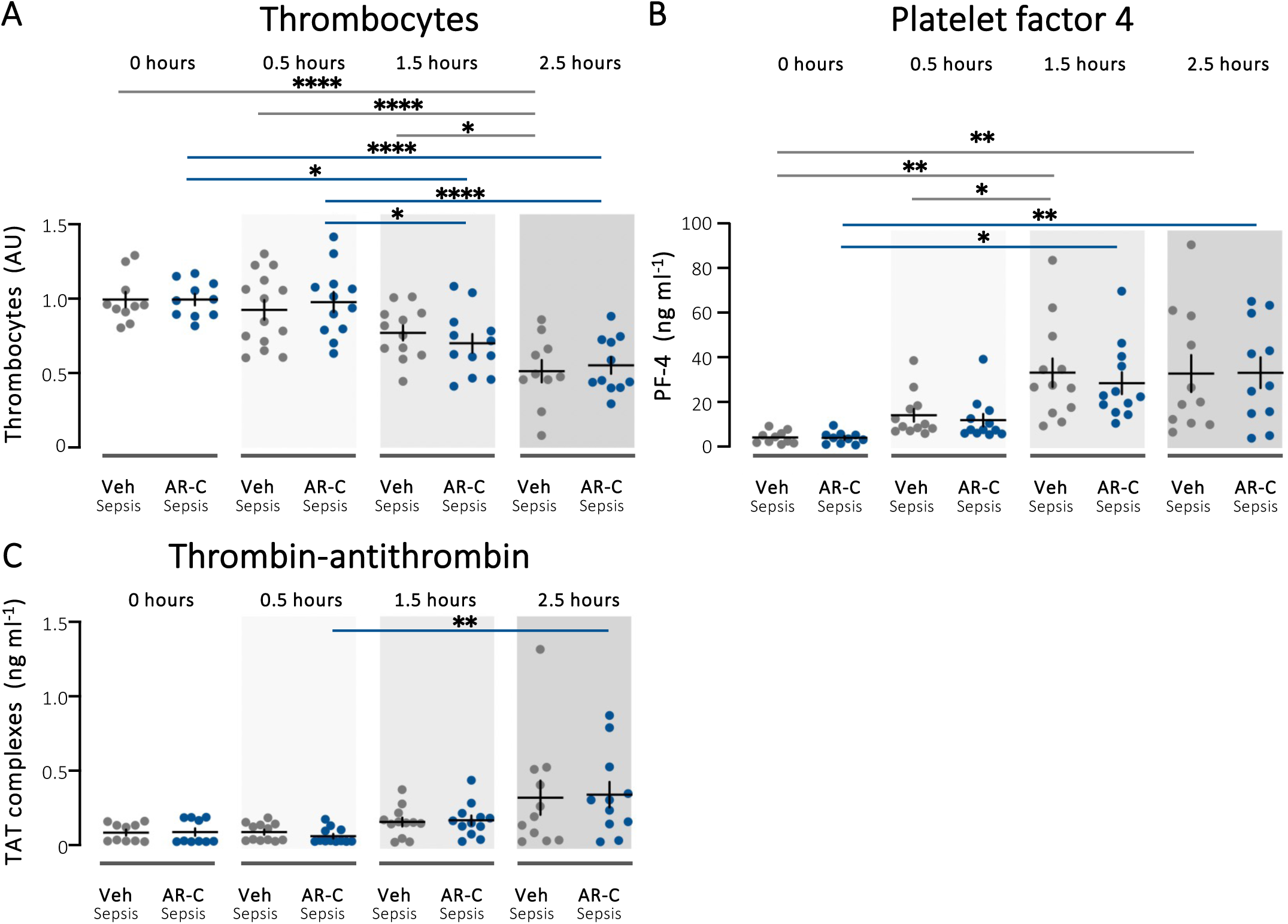
The sepsis-induced fall in circulating thrombocytes in WT mice infused with AR-C118925XX or vehicle. (**A**) shows the relative number of thrombocytes from 0-2.5 h after injection of 3.3 10^8^ E. coli (n=9-12 in the groups). (**B**) illustrate the corresponding change in thrombocyte activation marker platelet factor-4 (PF-4) in the same animals. (**C**) in reference, shows the change in intravascular formation of thrombin-antithrombin (TAT) complexes in plasma in the same animals as in A. Data are given as mean±SEM, * indicates p<0.05, **p<0.01, ***p<0.001 and ****p<0.0001.

**Supplementary Figure 3.**
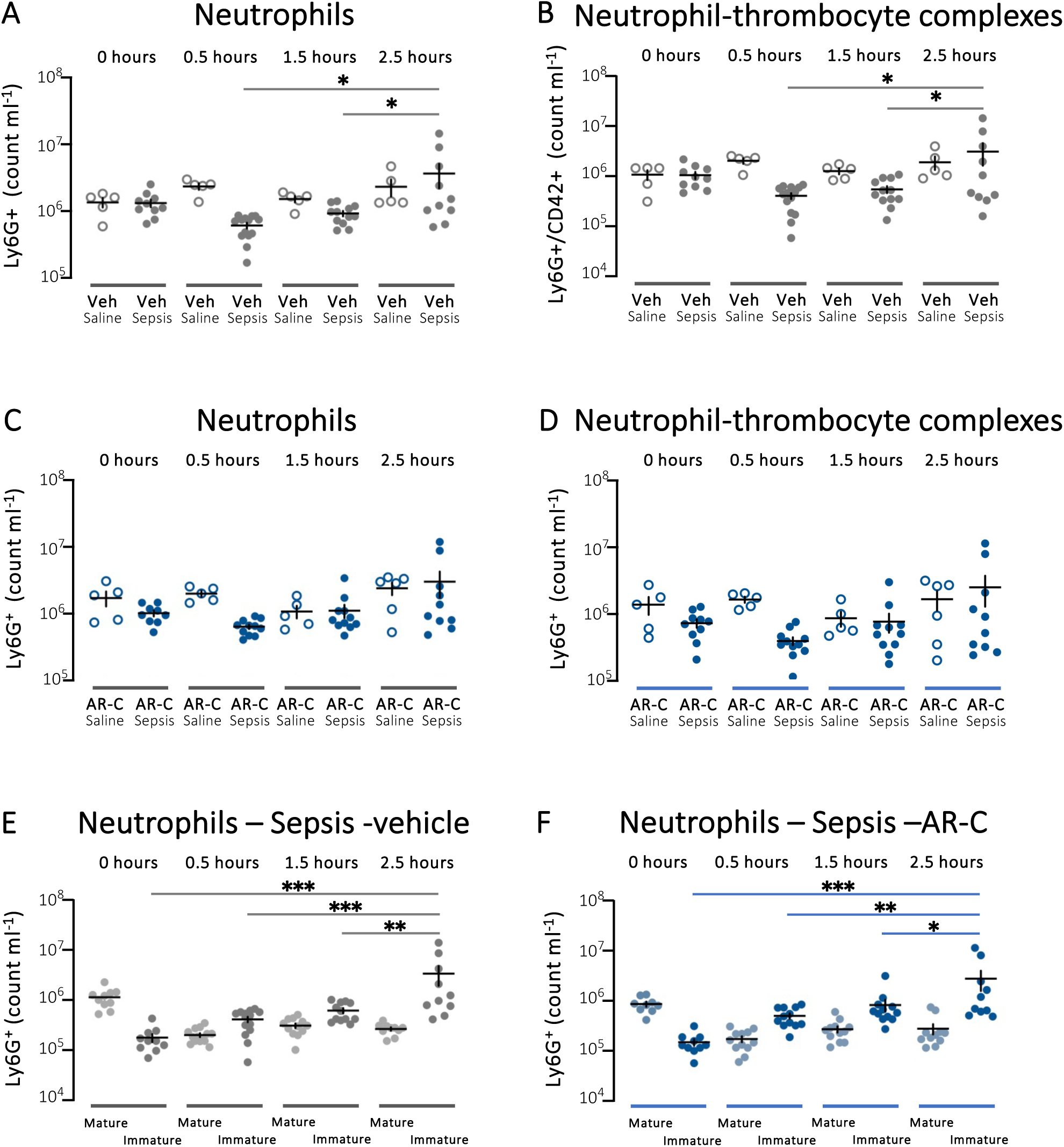
The sepsis-induced changes in circulating neutrophils and neutrophil-thrombocyte complexes in mice exposed to P2Y_2_ inhibitor or vehicle. (**A**) shows the neutrophil count in blood from WT mice infused with vehicle after injection of 3.3 10^8^ E. coli. (**B**) show the corresponding formation of neutrophil-platelet complexes. (**C**) shows the neutrophil count in blood from WT mice infused with AR-C118925XX (AR-C, 2.03 μg hour^-1^) after injection of 3.3 10^8^ E. coli. (**D**) show the corresponding formation of neutrophil-thrombocyte complexes. (**E**) illustrates the change in mature and immature neutrophils over time in WT mice infused with vehicle exposed to 3.3 10^8^ E. coli or saline, whereas (**F**) show the corresponding data in WT mice infused with AR-C118925XX. All data are given as mean±SEM, * indicates p<0.05, ** p<0.01, and ***p<0.001.

**Supplementary Figure 4.**
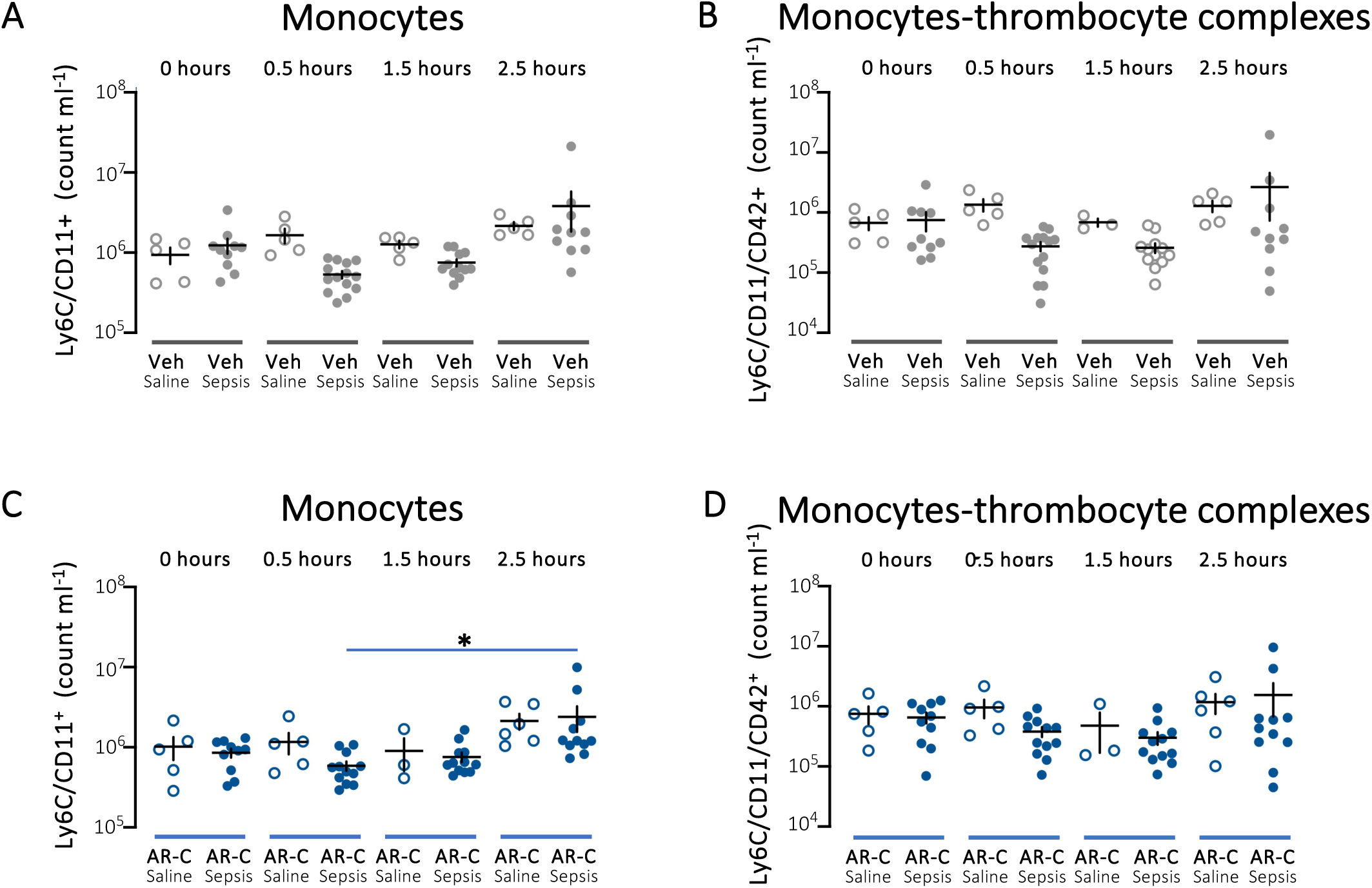
The sepsis-induced changes in circulating monocytes and monocyte-thrombocyte complexes in mice exposed to P2Y2 inhibitor or vehicle. (**A**) shows the monocyte count in blood from WT mice infused with vehicle after injection of 3.3 108 E. coli (**B**) show the corresponding formation of monocyte-thrombocyte complexes. (**C**) shows the monocyte count in blood from WT mice infused with AR-C118925XX (AR-C, 2.03 μg hour-1) after injection of 3.3 108 E. coli. (**D**) show the corresponding formation of monocyte-thrombocyte complexes. All data are given as mean±SEM, * indicates p<0.05.

**Supplementary Figure 5.**
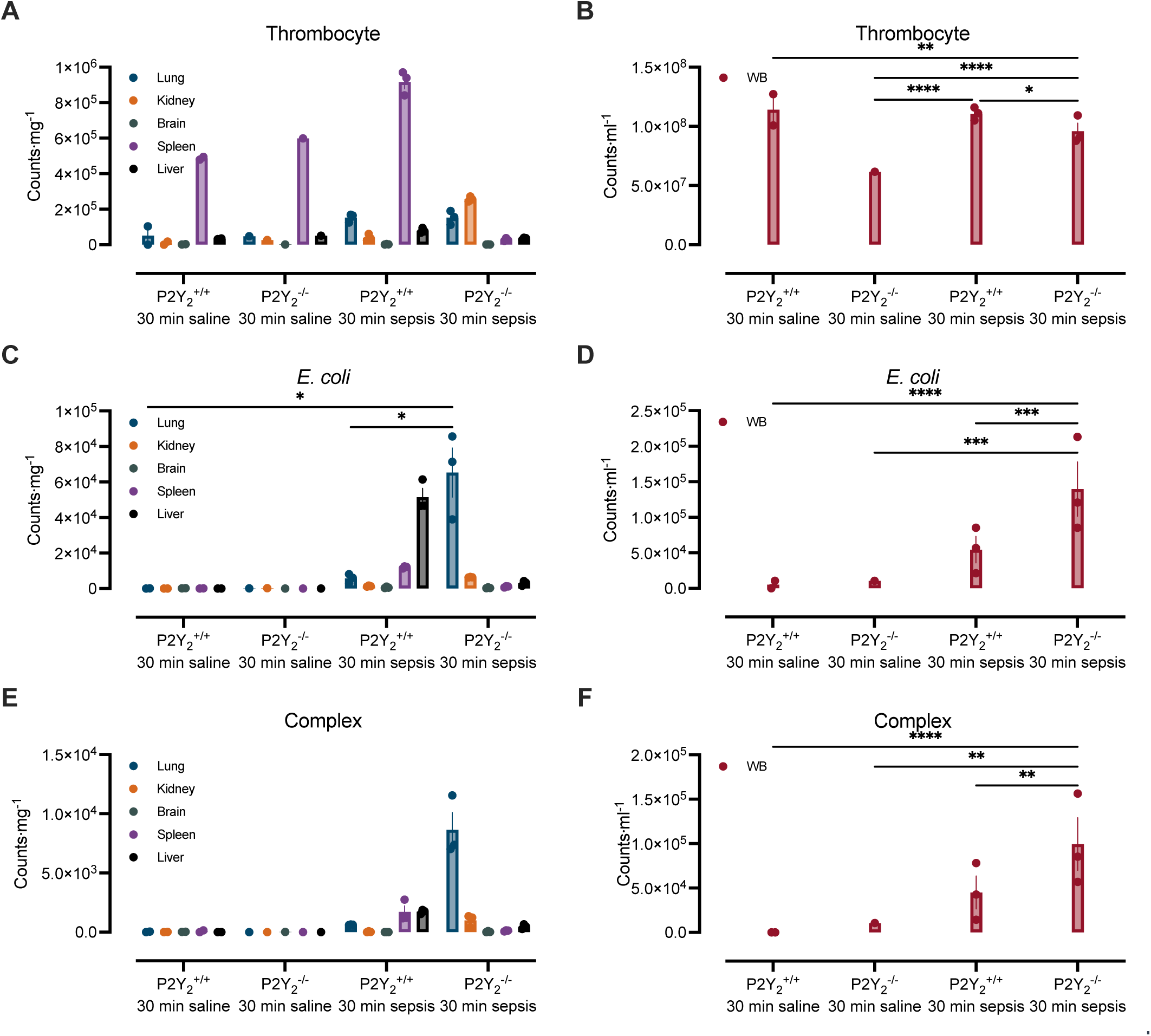
Shows the tissue-dependent clearance of UPEC in WT and P2Y ^-/-^ mice 30 minutes after injection of 3.3 10^8^ ARD-6/EGFP-pBAD. (**A**) illustrates the CD42^+^ counts per and the corresponding tissue accumulation of thrombocytes (**B**) and thrombocyte-UPEC complexes (**C**). Data are given per gram tissue, as mean±SEM, and * indicates p<0.05, ** p< 0.01, *** p<0.001 and ****p<0.0001.

